# An unmanned aerial vehicle pipeline to estimate body volume at scale for ecological monitoring

**DOI:** 10.1101/2023.11.23.567408

**Authors:** Thomas C Stone, Katrina J Davis

## Abstract

1. Demographic data are essential to construct mechanistic models to understand how populations change over time and in response to global threats like climate change. Existing demographic data are either lacking or insufficient for many species, particularly those that are challenging to study, such as marine mammals. A pipeline for collecting accurate demographic data to construct robust demographic models at scale would fill this knowledge gap for many species, including marine mammals like pinnipeds (seals, sea lions, and walruses).
2. We introduce a non-invasive pipeline to estimate the 3D body size (volume) of species that will allow monitoring at high spatial and temporal scales. Our pipeline integrates 3D structure-from-motion photogrammetry data collected via planned flight missions using off-the-shelf, multirotor unmanned aerial vehicles (UAVs). We apply and validate this pipeline on the grey seal *Halichoerus grypus*, a marine species that spends much of its time at sea but is predictably observable during its annual breeding season. We investigate the optimal ground sampling distance (GSD) for surveys by calculating the success rates and accuracy of volume estimates of individuals at different elevations.
3. We establish an optimal GSD of 0.8 cm px^-1^ for animals similar in size to UK grey seals (∼1.4 - 2.5 m length), making our pipeline reproducible and applicable to a broad range of organisms. Volume estimates were accurate and could be made for up to 68% of hauled-out seals in the study areas. Finally, we highlight six key traits that make a species well-suited to estimating body volume following this pipeline. Good candidates include large reptiles like crocodiles, large mammals such as hippopotamus, and shrubs or bushes in deserts and Mediterranean habitats.
4. Our pipeline accurately estimates individual body volume of marine macrovertebrates in a time-and cost-effective manner whilst minimising disturbance. Whilst the approach is applied to pinnipeds here, the pipeline is adaptable to many different taxa that are otherwise challenging to study. Our proposed approach therefore opens up previously inaccessible areas of the Tree of Life to demographic studies, which will improve our ability to protect and conserve these species into the future.

## Introduction

Population modelling has produced key insights into the ecology (Goldberg et al., 2001), evolution (Grant & Grant; 2002), and conservation biology (Baxter et al., 2006) of multiple species. Indeed, using demographic data, one can parameterise mechanistic models to understand how populations change through time (Thomas et al., 2019), evolve (Barfield et al., 2011), and can be managed in the most cost-effective manner (Yackulic et al., 2021; Taylor & Hastings, 2004). In this context, demographic data describe the size, structure, and/or trends through time of a population. These data are often used to parameterise structured population models (Easterling et al., 2000; Caswell 2001). Such models can describe how different components of a population contribute differently to the population growth rate through birth, survival, and migration (Sibly et al., 2002; Gerber & Heppell, 2004). Understanding these population changes is crucial given the increasing external threats populations face (Watson et al., 2019), and as such these changes are integral metrics of the IUCN conservation status of species (IUCN, 2022).

Robust estimates of population trends emerge from demographic models often parameterised with data possessing four qualities. These qualities are long-term (White 2019), individual-level (Merow et al., 2014), large sample size (Fiske et al., 2008), and, for species whose survival and reproduction are best described as a function of continuous variables (*e.g.*, size, height, weight), data on those continuous traits (Easterling et al., 2000; Ramula et al., 2009). Longer time series of demographic data can help us understand how populations change and the consistency of trends over time (White 2019). Individual-level data covering a high proportion of the population is important to capture the naturally occurring heterogeneity in vital rate values (*e.g.*, survival, growth, reproduction), a key aspect of population viability (Fiske et al., 2008). Continuous traits such as body size (as opposed to discrete traits that can only take certain values or states, *e.g.* developmental stage or age) are good predictors of vital rates in a vast number of species (Savage et al., 2004), as they tend to provide key insights into underlying mechanisms that shape fitness components. For example, body size is intrinsically linked to metabolic rate (Sparling et al., 2016) and competitive abilities (Whitman, 2008), which in turn shapes survival and reproduction (Brown et al., 2004; Savage et al., 2004). However, despite the preference for models based on data with these properties, we do not have such data for many species (Lebreton et al., 2012; Salguero-Gómez et al., 2015; Salguero-Gómez et al., 2016), even among highly charismatic ones (Conde et al., 2019).

High-quality demographic data are lacking for many animal species because they are not easily observed or approached (Cooke 2008; Sollmann et al., 2013), or because time-consuming and invasive methods such as capturing individuals or using sedatives are required (Manning & Goldberg, 2010; Hodgson et al., 2020; Williams et al., 2021). Consequently, we do not have high-resolution demographic data for species such as many marine species that are challenging to observe (Davis 2022). Examining data-deficient species is necessary to effectively protect them from threats, particularly as less understood species are often more threatened or face different challenges to better-studied species (Borgelt et al., 2022). For example, marine mammals have faced more threats than their better-understood terrestrial counterparts (Schipper et al., 2008). However, novel technologies such as UAVs (Unmanned Aerial Vehicles, drones) may help us overcome these challenges.

UAVs have diverse applications in ecology and conservation biology, ranging from crop monitoring to assessing forest health (Anderson & Gaston, 2013; Maimaitijiang et al., 2020; Sun et al., 2021; Ecke et al., 2022; Larsen & Johnston, 2023). Recent advancements have made UAVs suitable for collecting continuous state demographic data (*e.g.*, body size) from populations that would otherwise be challenging to study (Krause et al., 2017; Shero et al., 2021; Christiansen et al., 2022). Indeed, UAVs can access remote areas, eliminating the need to approach, sedate, and handle dangerous or threatened species (Krause et. al., 2017; Shero et al., 2021). Disturbance to study organisms can also be minimised by considering species-specific hearing sensitivity and operating UAVs at higher altitudes without sacrificing on high resolution data (Duporge et al., 2021). Using UAVs is cheaper and quicker than traditional monitoring methods (Jackson et al., 2022) and can be applied over large spatial scales (Bogdan et al., 2021). To date, though, UAVs have been used for 2D size estimates of individual animals, including at the population scale (Gray et al., 2019; de Kock et al., 2021; Infantes et al., 2022). However, 3D body size (volume) estimates can also be estimated using UAVs by structure-from-motion photogrammetry (Westoby et al., 2012), taking a series of overlapping images from a known altitude that are stitched to create a geometrically accurate 3D model (Hodgson et al., 2020; Shero et al., 2021). Recently, multiple studies have shown that 3D body size (body volume) estimates from UAVs are more accurate than 2D estimates, as the former are less sensitive to body position (Hodgson et al. 2020; Shero et al., 2021). Despite this advancement, current methods estimate 3D body volume of a single individual or small group of individuals at a time, and so are not easily scalable to whole populations (Hodgson et al., 2020; Shero et al., 2021). Therefore, there is need for a pipeline to estimate body volume at the population scale.

Here, we develop and validate a pipeline to accurately estimate body volume at scale using commercially-available and broadly-affordable (<£6,000) UAVs. We demonstrate our approach on the grey seal *Halichoerus grypus*, a cryptic marine species that spends much of its time at sea but is predictably observable annually during its breeding season when it hauls-out on land (Hall & Russell, 2018). We conducted fieldwork on two regionally-important grey seal populations in the UK: the Isles of Scilly and the Farne Islands. Our pipeline obtains accurate body volume estimates for 68% of hauled-out individuals at the identified optimal ground sampling distance (GSD) of 0.8 cm px^-1^. We establish minimum criteria to determine if different UAVs can be used for estimating body volumes. Finally, we critically evaluate the suitability of a range of species for body volume estimates by UAV and highlight the key characteristics that make a species more suitable for these methods. Thus, we demonstrate that our pipeline is applicable for a broad range of taxa, opening areas of the tree of life for demographic modelling that were previously inaccessible.

## Methods

### Study Species

The grey seal *Halichoerus grypus* is a long-lived (20+ years for males, 30+ years for females; SCOS 2021) marine apex predator species that spends most of its time at sea (McConnell et al., 1999). Grey seals haul-out on land to rest and in larger colonies to pup and moult (Hall & Russell, 2018), so some of their key life history traits are predictably observable annually. The UK is home to ∼35% of grey seals worldwide (SCOS, 2021), and UK populations are increasing in size from historic lows due to hunting (Lambert, 2002; Russell et al., 2019). Grey seals have a polygynous mating system and mature females can give birth to one pup per year that is weaned at approximately three weeks (Hall & Russell, 2018). The pup then moults its white coat to gain adult pelage and after another three weeks leaves the colony (Hall & Russell, 2018). In this study, we assigned any individual with a white coat to a ‘pup’ category, and any individual with adult pelage to an ‘adult’ category.

### Study Sites

Grey seal populations on the Isles of Scilly and Farne Islands (Fig. 1A) both make important contributions to regional grey seal pup production (SCOS 2021). Approximately 50% of Southwest England’s grey seal pup production takes place in the Isles of Scilly, which on average is inhabited by ∼600 individuals (Sayer & Witt, 2018). The Farne Islands’ grey seal population produces ∼3,000 pups annually, accounting for ∼25% of England’s grey seal pup production (SCOS, 2021). Access to these sites is challenging, particularly in the Isles of Scilly, due to a lack of suitable boat landing spots and prevailing sea and weather conditions. Therefore, surveys are largely undertaken from sea (Sayer & Witt, 2018), with the use of visual aids such as binoculars. Visual surveys from sea will necessarily fail to spot seals hidden from view or located in the centre of islands, potentially leading to inaccuracies in counts.

**Figure 1.**
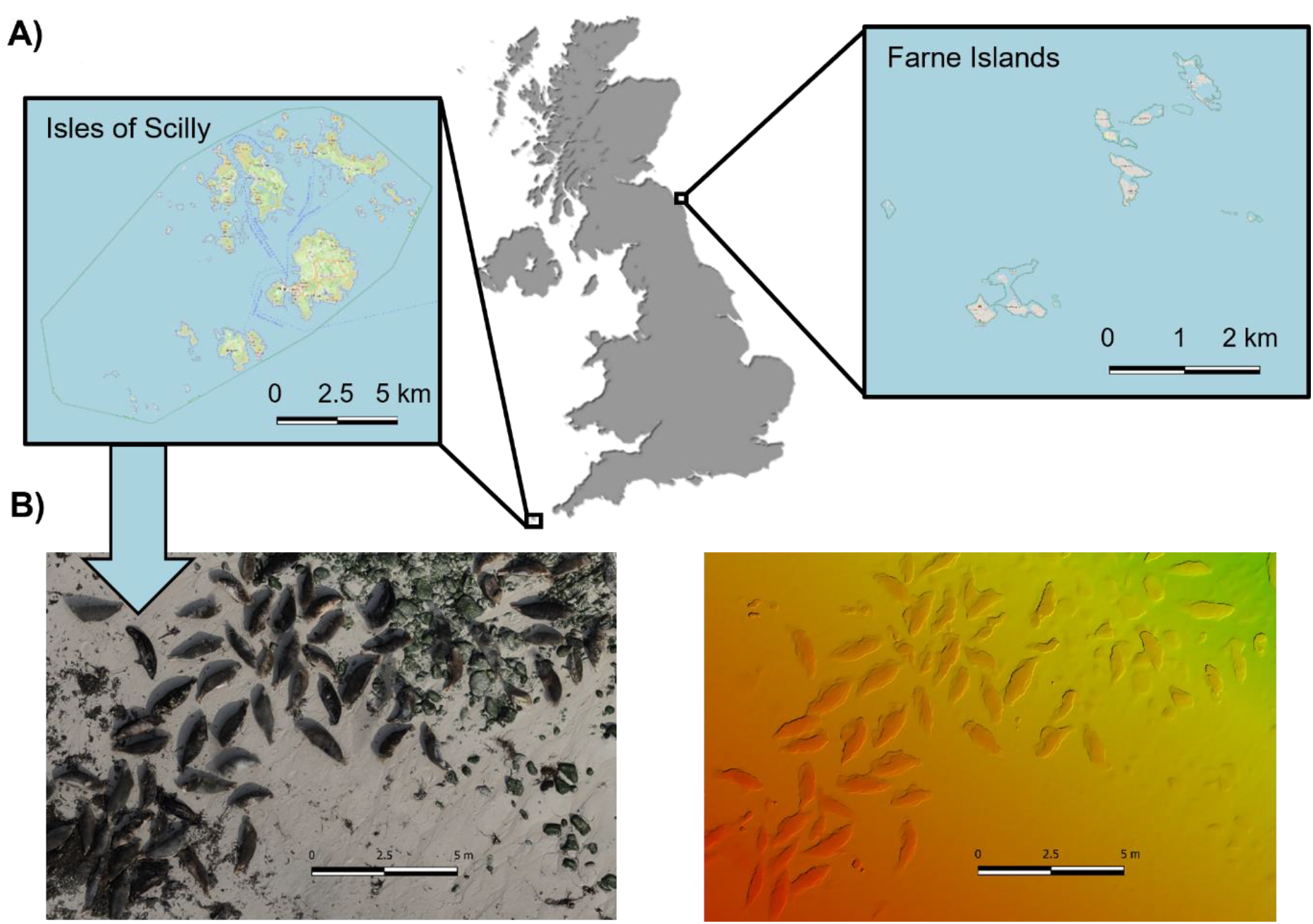
The locations of the two field sites used to test our pipeline for estimating body volume using grey seals *Halichoerus grypus* as a case study, together with an example of what an orthomosaic (geometrically accurate 2D representation of the survey area) and a digital surface model (DSM, 3D representation of the survey area showing the elevation of objects above the Earth’s surface) obtained following our pipeline look like. **A**: A map showing the location of our two field sites in the United Kingdom. The Isles of Scilly (left) consist of five inhabited and > 100 uninhabited islands ∼ 45 km off Southwest England. The Farne Islands (right) consist of 15 - 20 (tide-dependent) small islands ∼ 2 – 10 km off Northeast England, uninhabited except for a ranger station on one island (Inner Farne). **B**: Left: An orthomosaic generated from unmanned aerial vehicle (UAV) images taken from an island in the Isles of Scilly and Right: the corresponding digital surface model, with red colours indicating higher elevation and green colours indicating lower elevation.

### Field Technologies

We used two different off-the-shelf, multirotor UAVs to carry out survey missions^1^. These UAVs were a DJI Mavic 2 Enterprise Advanced (DJI; Shenzhen, China; unfolded diagonal wingspan 354mm, take-off weight 909 g, GPS accurate to ± 0.5 m vertically and ± 1.5 m horizontally, £5,480) and a DJI Mavic 2 Pro (unfolded diagonal wingspan 354 mm, take-off weight 907 g, GPS accurate to ± 0.5 m vertically and ± 1.5 m horizontally, £1,349). The visual-range camera on the DJI Mavic 2 Enterprise Advanced has a minimum ground sampling distance (GSD, a measure of image quality; Box 1) 20% lower than the DJI Mavic 2 Pro, meaning the DJI Mavic 2 Enterprise Advanced takes higher resolution images. The DJI Mavic 2 Enterprise Advanced can also take infrared images, which can be useful for identifying seals as they appear similar to rocks on RGB images but have an active metabolism that enables them to be more easily identified from thermal imagery.

### Survey Planning & Flight Parameters

To determine which islands to survey, we consulted site managers and conducted scouting missions in the field. We pre-programed survey missions in DJI Pilot (DJI; Shenzhen, China. v2.5.1.15) or Fly Litchi (VC Technology; London, United Kingdom. V4.25.0-a) following transect routes to take a series of overlapping photos across a seal colony with the gimbal angled 90 degrees vertically down. This transect pattern is a key difference to previous pinniped body volume work that either used radial flight patterns circling a small group of seals (Shero et al., 2021), or focussed on a single individual at a time (Hodgson et al., 2020). Our transect method enables a wider area to be covered with a single survey. Early trials showed no difference in accuracy between volume estimates from transect or radial flight patterns. Sufficient overlap between images is required so images can be stitched together to create an orthomosaic (geometrically accurate 2D representation of the survey area, Fig. 1B) and a digital surface model (DSM, 3D representation of the survey area showing elevation of objects above the Earth’s surface, Fig. 1B). We set frontal image overlap at 80% and side overlap at 70%, meaning 80% of the previous image was in the next image and 70% of the area covered in the previous transect was in the next transect. Transect routes can also be saved and so are reproducible across years.

### Data Collection

Ideally, censuses should be timed to maximise the number of individuals that can be captured within the survey, and when key vital rates are measurable (*e.g.*, peak reproduction, juvenile to adult transition, *etc.*). For grey seals, surveying during the pupping season when much of the population are hauled-out on land maximises the number of individuals captured in surveys (Sayer and Witt, 2018). In particular, breeding female grey seals and their pups can be captured in surveys during the pupping season. In the Isles of Scilly, the pupping season lasts from September to November (Sayer and Witt, 2018). We censused the Isles of Scilly grey seal population by UAV in late-September 2022 over four days, covering every island with > five individuals based on previous surveys (Sayer et al., 2012; Sayer & Witt, 2018). In total, we surveyed 21 islands over 23 individual missions in the Isles of Scilly. The Farne Islands’ pupping season lasts from November to December (Coulson & Hickling, 1964; SCOS, 2021). We censused every island in the Farne Islands with suspected seal presence (14 islands over 15 individual missions), as informed by local rangers.

For our UAV surveys of grey seals, we launched a combination of survey missions from land (*n* = 12) and from a small boat (*n* = 26). Launching missions from boats minimises disturbance to seals and precludes the need to make difficult landings on islands when conditions are poor. By contrast, launching UAV missions from land is generally safer for operators, quicker, and easier in poor weather conditions, allowing longer flight times since less battery reserves are needed for landing. We launched missions from land or boat depending on the accessibility of landing sites at a safe distance from seals. We maintained an in-flight altitude of ≥ 40 m when surveying and took-off and landed UAVs > 50 m from seals to avoid disturbance (Pomeroy et al., 2015; Larsen & Johnston, 2023). Seal and seabird behaviour were monitored during flights to ensure no disturbance. During the surveys, no abnormal seal behaviour was observed that could be attributed to our UAV missions.

### Data Processing

To estimate body volume on a population-scale, we aimed to find the most efficient way to process the image data we collected. This task involved establishing the optimal way to conduct pre-processing on the images collected, construct 3D models from these images, locate seals in the model, and estimate their body volumes. Before processing the images collected, we removed blurry images (< 5% in our case) and ensured auto-collected image metadata including altitude and GPS coordinates were correct. We constructed 3D representations of the survey area using structure-from-motion photogrammetry with the software ‘Pix4D Mapper’ (v. 4.7.5; Pix4D S.A., Prilly, Switzerland). As a series of overlapping photos were taken at known altitudes, every point in the survey area is present in many images from many angles meaning a geometrically accurate 3D model can be reconstructed from the 2D images taken using structure-from-motion photogrammetry (Hodgson et al., 2020; Shero et al., 2021). We used the default settings of the ‘3D Maps’ Processing Option in Pix4D, except for changing the minimum number of matched keypoints (points Pix4D can easily recognise and match between multiple images) from three to five to reduce sensitivity to seal movements between photos (Shero et al., 2021). We processed projects in Pix4D Mapper to create an orthomosaic (Fig. 1B) and a digital surface model (DSM, Fig. 1B) for each discrete area we surveyed. We manually located each seal within each orthomosaic. Any seals in the water were counted but their volumes were not estimated as it is currently only possible to estimate the volume of seals on land (but alternative methods and techniques have emerged that may contribute in this context: Chirayath & Earle, 2016; Chirayath & Li, 2019; Hirtle et al., 2022).

Next, we estimated the volume of each seal in Pix4D Mapper using the Volume function by manually marking vertices around a seal. The base of each individual seal was constructed using the ‘Triangulated Plane’ setting by triangulation of all marked vertices. Pix4D calculates volume as the difference in altitude between the base and the top of the seal using the DSM. We estimated the volume of 196 individual adult seals (out of a possible 437) and 48 pups (out of a possible 183) from 21 islands in the Isles of Scilly and one island (Inner Farne) in the Farne Islands (Table 1). The accuracy of volume estimates was tested by applying our pipeline to a large, inflatable object of known volume (see later section ‘Verification of accuracy’). We compared sizes between adult grey seals from the Isles of Scilly and the Farne Islands.

**Table 1.** The success rate of reconstructing grey seals *Halichoerus grypus* in 3D so their volumes could be estimated following our pipeline was 2.5 times higher at smaller ground sampling distances (GSDs, Box 1) and overall 1.7 times higher for adults compared to pups. Volume estimates were attempted at the different GSDs listed below, along with the altitude corresponding to these GSDs with the two different UAVs used for surveys. The number of grey seal pups and adults whose volume we attempted to estimate, and the percentage success rates of estimating their volumes, at the different GSDs we tested. The altitude at which the DJI Mavic 2 Enterprise Advanced achieves a GSD of 1.4 cm px^-1^ and the DJI Mavic 2 Pro achieves a GSD of 0.8 cm px^-1^ are listed as ‘N/A’ because we did not carry out any surveys with these UAV at these GSDs. The rate of successfully estimating adult seal volumes was significantly higher for adults compared to pups at GSDs of 0.8cm px^-1^ and 1.2 cm px^-1^.

| **GSD**<br>**(cm**<br>**px<sup>-1</sup>)** | **DJI Mavic 2**<br>**Enterprise**<br>**Advanced**<br>**Altitude (m)** | **DJI Mavic 2**<br>**Pro**<br>**Altitude (m)** | ***n* Adults** | **%**<br>**Success**<br>**Adults** | ***n* Pups** | **%**<br>**Success**<br>**Pups** | ***P* Value**<br>**Adult Vs**<br>**Pup**<br>**Success** |
| --- | --- | --- | --- | --- | --- | --- | --- |
| 0.8 | 40 | N/A | 154 | 68 | 88 | 41 | < 0.001 |
| 1.0 | 50 | 40 | 178 | 27 | 27 | 11 | 0.12 |
| 1.2 | 60 | 50 | 87 | 28 | 59 | 10 | 0.019 |
| 1.4 | N/A | 60 | 18 | 44 | 9 | 0 | NA |

### Determining Optimal Ground Sampling Distance

Most fieldwork activities typically operate under a delicate data quality *vs.* cost trade-off, whether costs be time and/or financial (McDonald-Madden et al., 2010). Here, the key components of data quality are the percentage of individuals that could be reconstructed within Pix4D so their volumes could be estimated and the accuracy of those volume estimates. Both qualities are dependent on ground sampling distance (GSD, Box 1), which is optimised with lower flight altitudes or a UAV that takes higher resolution images. However, UAVs that take higher resolution images are more expensive (*e.g.*, DJI M300 RTK with DJI Zenmuse P1 camera, £13,330; compared to the DJI Mavic 2 Enterprise Advanced, £5,480). Conducting missions at lower altitudes means surveying takes longer as less area is captured per image. This trade-off between data quality and cost is highlighted by the difference in total mission time for the Isles of Scilly census. If all survey missions were planned to be undertaken at a minimum GSD of 0.8 cm px^-1^ (40 m altitude with the DJI Mavic 2 Enterprise Advanced), the time required to be spent in the field would increase by 35% compared to surveying at a GSD of 1.2 cm px^-1^ (60 m altitude, see Appendix A for further details). Extra time in the field incurs higher financial costs. Overall, the highest GSD at which volume estimates are accurate and a high percentage of animal volumes can be estimated gives the optimal GSD to undertake surveys whilst minimising the cost component of the trade-off.

### Success of Estimating Seal Volumes

To help determine the optimal minimum GSD for resolving the data quality against cost trade-off, we calculated the percentage of seals that could be successfully reconstructed in Pix4D to obtain volume estimates at different GSDs (Fig. 2). We tested GSDs of 0.8 cm px^-1^ (40 m altitude with DJ Mavic 2 Enterprise Advanced; *n* = 154 adults, 88 pups), 1.0 cm px^-1^ (50 m; *n* = 178 adults, 27 pups), 1.2 cm px^-1^ (60 m; *n* = 87 adults, 59 pups), and 1.4 cm px^-1^ (70 m; *n* = 18 adults, 9 pups) (Table 1). We did not test altitudes lower than 40 m to avoid disturbing seals (Pomeroy et al., 2015; Larsen & Johnston, 2023). All 23 missions from the Isles of Scilly and one mission from the Farne Islands were analysed here. We introduced the mission from the Farne Islands to balance the sample size at a GSD of 0.8 cm px^-1^. We assessed whether there were significant differences between the percentage of volume estimates possible for adult and pup grey seals at GSDs of 0.8 cm px^-1^, 1.0 cm px^-1^ and 1.2 cm px^-1^. We expected the success rate for reconstructing seals would be higher at lower GSDs with higher resolution images. We also expected success rates would be higher for adults than pups because pups are typically five-to tenfold smaller than adults (Hall & Russell, 2018) and so more difficult for Pix4D to reconstruct accurately. The sample size is lower at a GSD of 1.4 cm px^-1^ because accuracy tests showed that even at a GSD of 1.2 cm px^-1^, the resolution of images was insufficient to accurately estimate volumes (see ‘Verification of Accuracy’ section). We therefore did not carry out any statistical tests at a GSD of 1.4 cm px^-1^, although we still attempted to estimate the volumes of seals at this GSD for completeness.

**Figure 2.**
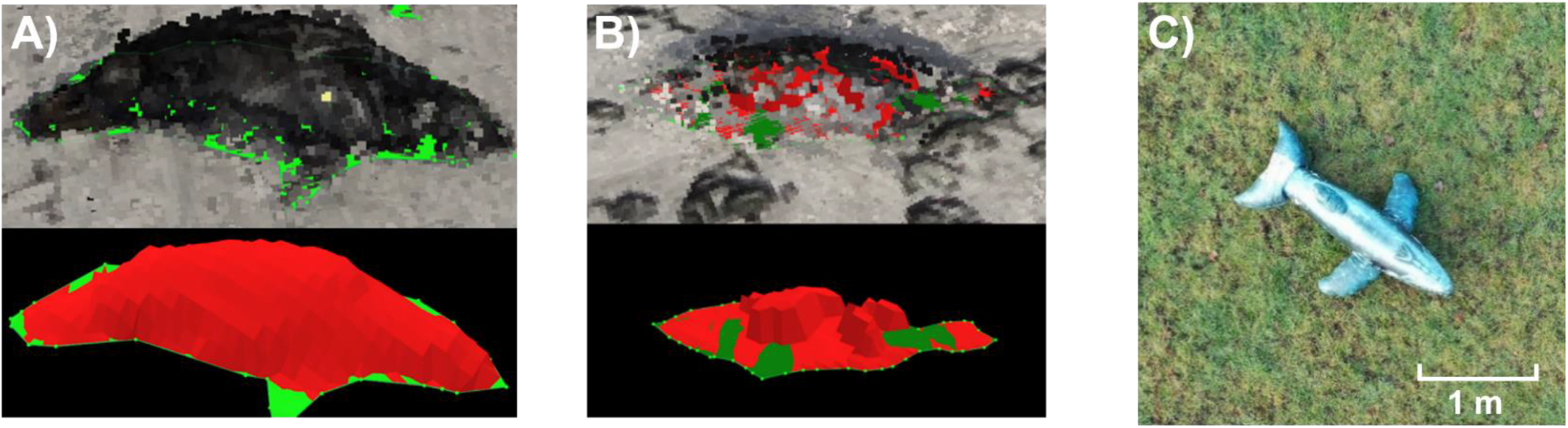
To establish the optimal ground sampling distance (GSD, Box 1) at which to carry out surveys of grey seals *Halichoerus grypus* to estimate their body volumes, we calculated the success rates of being to estimate seal volumes and the accuracies of volume estimates at different GSDs. We used the software Pix4D Mapper to reconstruct seals from UAV photographic data so that their volumes could be measured. Here, we compare a seal that has been successfully reconstructed so its volume can be measured to a seal where the programmatic reconstruction failed. As validation, we also used an inflatable object of known volume to test the accuracy of volume estimates made following our pipeline. **A:** A successfully reconstructed seal that has had its volume estimated. The images used had a GSD of 0.8 cm px^-1^. Note that the upper image appears pixilated because the step of marking the base of the object to estimate the volume of objects is performed in the dense point cloud. **B:** A seal that has not been reconstructed successfully, likely because it moved between images in the survey. The images used had a GSD of 0.8 cm px^-1^. The volume of this seal could not be estimated. **C:** Unmanned Aerial Vehicle (UAV) image of a large inflatable object of known volume, which we used to test the accuracy of volume estimates obtained following the pipeline we developed.

### Verification of Accuracy

We tested the accuracy of volume estimates from UAVs using a large, inflatable, plastic object of known volume with a comparable size and shape to that of an adult seal (Fig. 2C. Note: a realistic inflatable seal was not available) following the same survey and processing pipeline used in the field. We carried out 30 UAV surveys of the inflatable with the DJI Mavic 2 Enterprise Advanced, 10 at each planned GSD of 0.8 cm px^-1^ (40 m), 1.0 cm px^-1^ (50 m) and 1.2 cm px^-1^ (60 m). We compared the mean volume estimated by UAV to the true volume of the object to assess the accuracy of volume estimates.

## Results

### Body Size Estimates

We calculated the mean body volume of grey seal adults and white coat pups from the Isles of Scilly and Inner Farne, Farne Islands using the Volume function in Pix4D Mapper (Fig. 3). The mean body volume in the Isles of Scilly for adults seals was 132.5 L ± 47.3 SD (*n* = 152) and 23.1 L ± 6.1 for pups (*n* = 9). From Inner Farne, the mean body volume of adult seals was 134.6 L ± 48.9 (*n* = 44) and 22.4 L ± 12.3 for pups (*n* = 39). A two-sample t-test showed no difference in the mean volumes of adult grey seals between the Isles of Scilly and Inner Farne (*pdf = 68.1* = 0.799). However, we note that the sample size in Inner Farne (*n* = 44) was smaller than the Isles of Scilly (*n* = 152), and that Inner Farne is not representative of the whole Farne Islands seal population because the Inner Farne haul-out consists almost entirely of females and pups.

**Figure 3.**
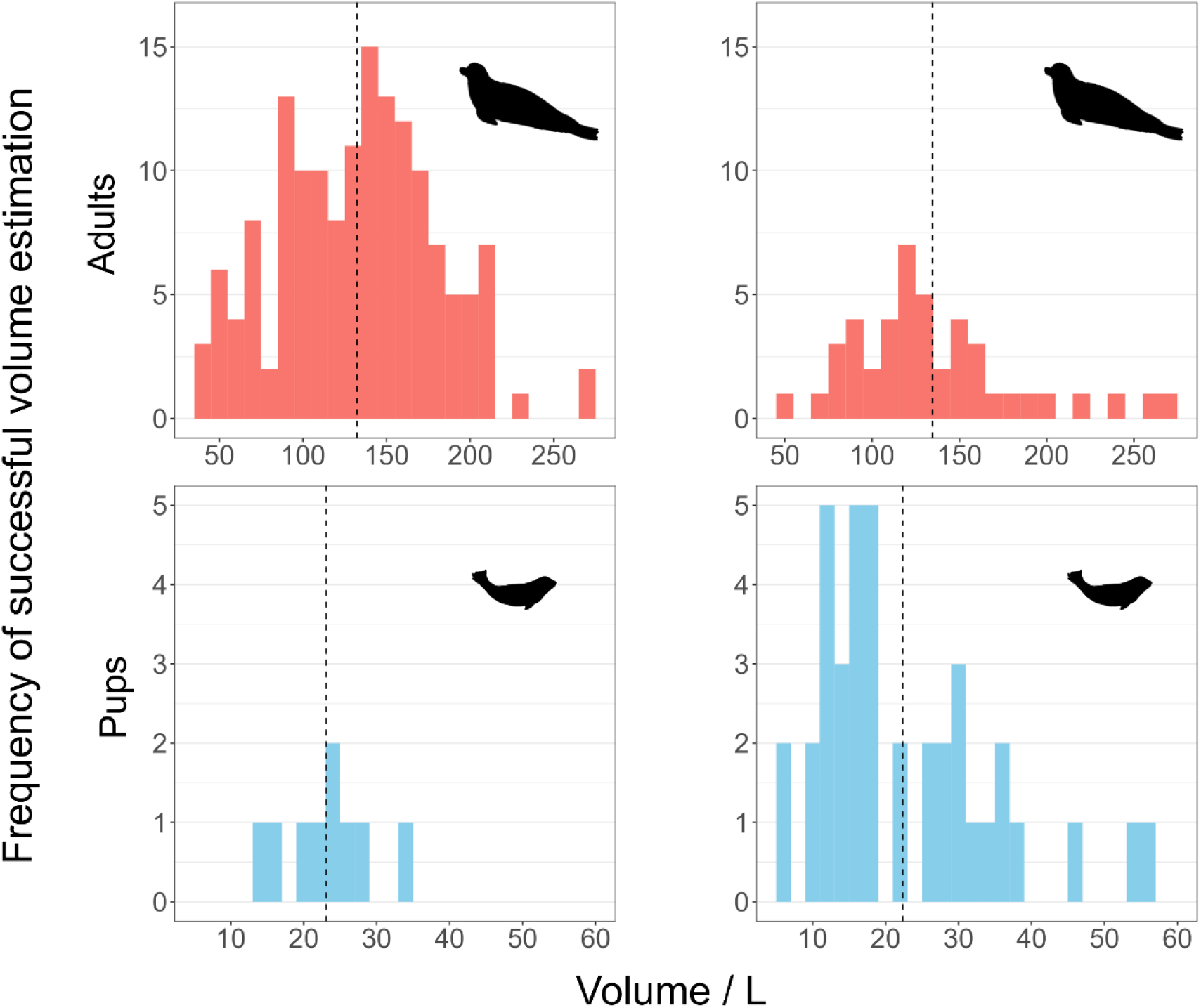
We estimated the body volume of grey seals *Halichoerus grypus* in the Isles of Scilly (*n* = 161) and Inner Farne, the Farne Islands (*n* = 83) using 3D structure-from-motion photogrammetry from UAV imagery. The histograms show the variation in body volume estimated by UAV following our pipeline, in Litres, for adult and white-coat pup grey seals in each location. Seals were assigned either to adult or pup categories depending if they had moulted the white coat associated with pups or not, meaning the adult category includes juvenile individuals. The dashed lines represent the mean volumes. The top two panels correspond to adult seals in (A) the Isles of Scilly, and (B) the Farne Islands, while the two bottom panels correspond to pups in (C) the Isles of Scilly and (D) the Farne Islands.

### Determining Optimal Ground Sampling Distance

To test the accuracy and applicability of our pipeline to estimate body volume on a population-scale, we calculated A) the percentage success rate of reconstructing a seal so its volume could be estimated (Fig. 2), and B) the accuracy of volume estimates, at different ground sampling distances (GSDs; Box 1).

#### A) Success of Seal Reconstruction

The percentage success rate of reconstructing individual seals decreased as GSD increased (Fig. 4A; Table 1). At a GSD of 0.8 cm px^-1^, the volume of 68% of adult seals (*n* = 154) and 41% of pups (*n* = 88) could be correctly estimated. The success rate decreased to 27% for adults (*n* = 178) and 11% for pups (*n* = 27) at 1.0 cm px^-1^, and was 28% for adults (*n* = 87) and 10% for pups (*n* = 59) at 1.2 cm px^-1^. The sample size for a GSD of 1.4 cm px^-1^ is low at 18 adults and nine pups. Of these, the volumes of 44% of adults and 0% of pups were estimated successfully. Successful estimation of volumes was much more likely for adults than pups at GSDs of 0.8 cm px-1 (*pdf = 1* = 0.0000624, *n* = 242) and 1.2 cm px^-1^ (*pdf = 1* = 0.0189, *n* = 146; table 1). At a GSD of 1.0 cm px^-1^ volume estimates were equally unsuccessful for adults and pups (*pdf = 1* = 0.124, *n* = 205), although the sample size for pups was relatively low at this GSD. Given the camera resolution of the UAVs used here, a GSD of 0.8 cm px^-1^ is optimal for surveying animals the size of UK grey seals (typically ∼1.4 - 2.5 m). This GSD achieves an optimal trade-off between data quality and costs, despite the longer time required to survey at this lower GSD (see ‘Determining Optimal Ground Sampling Distance’). Alternatively, if time is the limiting factor, a more expensive UAV (*e.g.*, DJI M300 RTK with DJI Zenmuse P1 camera, £13,330) that can survey at the same GSD from a higher altitude could be purchased.

**Figure 4.**
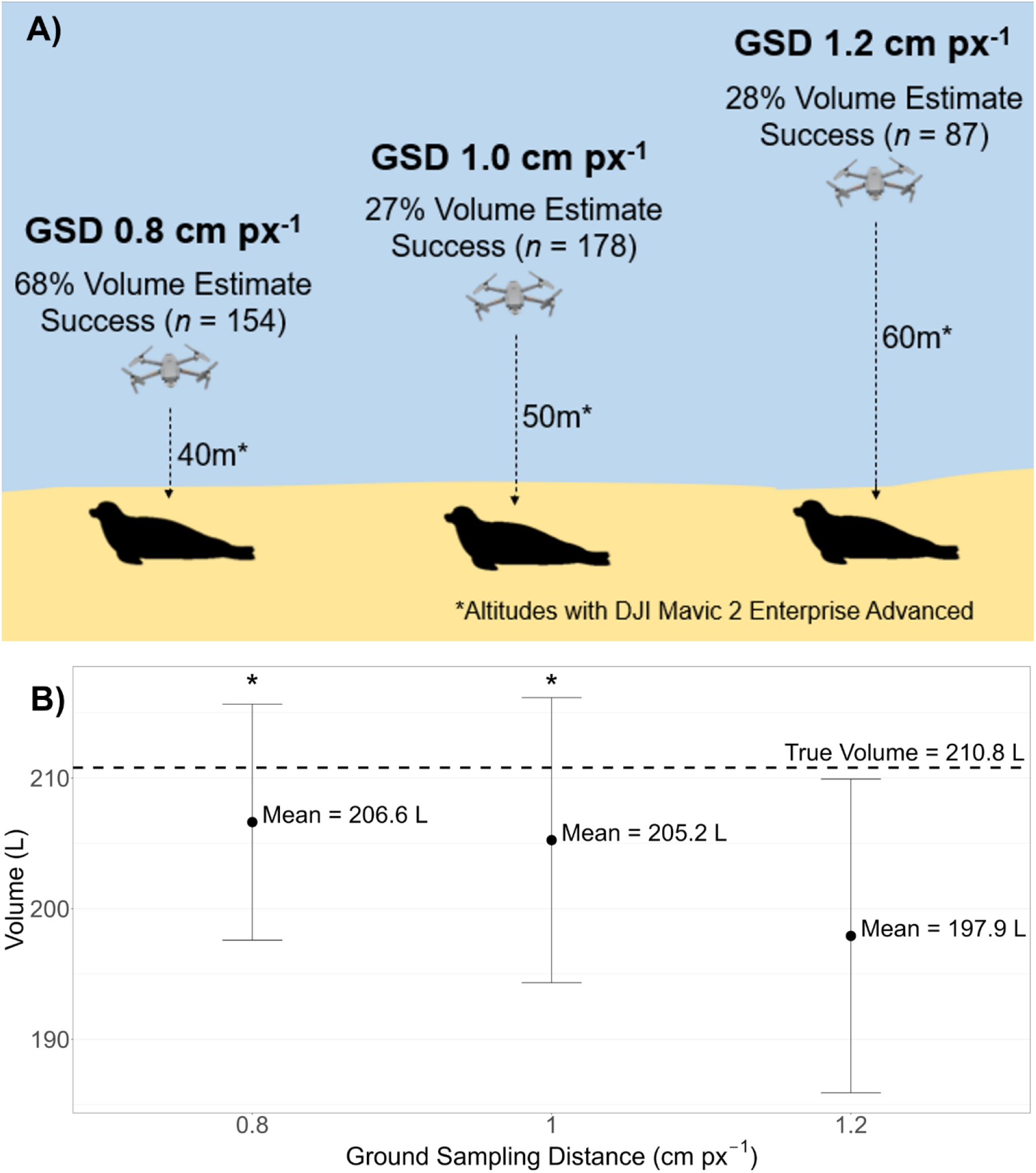
The optimal ground sampling distance (GSD; Box 1) at which to carry out surveys was 0.8 cm px^-1^, from the GSDs we tested. This conclusion was reached because the highest success rate of estimating seal volumes was at a GSD of 0.8 cm px^-1^, and because volume estimates are accurate at this GSD. **A:** A visualisation of the different percentage success rates at reconstructing seals within Pix4D Mapper so their volumes can be estimated at the different tested ground sampling distances of 0.8, 1.0 and 1.2 cm px^-1^. The highest success rate of 68% was obtained at a GSD of 0.8 cm px^-1^, followed by 27% at 1.0 cm px^-1^ and 28% at 1.2 cm px^-1^. Although we carried out some surveys at a GSD of 1.4 cm px^-1^, this GSD is not represented here because of a low sample size. **B:** We tested the accuracy of volume estimates obtained following our pipeline at different GSDs by comparing the mean volume estimates obtained to the true volume of a large, inflatable object using a one-sample t-test. The mean and standard deviation of volume estimates at the different tested ground sampling distances of 0.8, 1.0 and 1.2 cm px^-1^ are shown here. *: volume estimates at that ground sampling distance are not significantly different (p < 0.05) from the true volume, 210.8 L, represented with the dashed horizontal line.

#### B) Verification of Accuracy

We tested the accuracy of volume estimates by comparing estimates obtained following our pipeline to the true volume (210.8 L) of a large inflatable object (Fig. 4B). A one-sample t-test showed there was no statistically significant difference between the mean volume estimates obtained by UAV and the true volume of the inflatable at GSDs of 0.8 cm px^-1^ (*p* = 0.178, *n* = 10) and 1.0 cm px^-1^ (*p* = 0.142, *n* = 10). However, at 1.2 cm px^-1^, the mean volume estimated by UAV was significantly different to the true volume of the inflatable (*p* = 0.008, *n* = 10). Therefore, estimates of volume accuracy decrease with increasing GSD and thus altitude, as expected. In addition, the standard deviation of volume estimates increased from 9.04 L at 0.8 cm px^-1^ to 10.91 L at 1.0 cm px^-1^ and 12.01 L at 1.2 cm px^-1^. This result was expected, as lower resolution images should lead to more variable volume estimates. Based-on these results, we conclude that grey seal volume estimates obtained following our pipeline are accurate at GSDs of 0.8 cm px^-1^ and 1.0 cm px^-1^, whilst accuracy decreases with GSD.

## Discussion and Conclusions

We introduce a pipeline to estimate the body volume of animals > 80 L in size at a population-scale in a time-and cost-effective manner whilst minimising disturbance, using grey seals as case study species. We show that body volume estimates are accurate at ground sampling distances (GSDs, Box 1) achievable with off-the-shelf, multirotor unmanned aerial vehicles (UAVs) operated at altitudes that do not cause disturbance. Moreover, we establish that planning UAV surveys at a GSD of 0.8 cm px^-1^ optimises the trade-off between data quality and costs when estimating body volume. Our pipeline is highly adaptable to other species that possess similar traits to grey seals. The ability to obtain body volume estimates at the population-scale for a range of different species opens many possibilities for new research, including parameterising demographic models to help fill the demographic data gaps that exist across the tree of life (Lebreton et al., 2012; Salguero-Gómez et al., 2015; Salguero-Gómez et al., 2016).

Our pipeline enables individual body volumes to be estimated at the population scale. Figure B2 (Box 2) presents an overview of our pipeline, and the five stages of the pipeline are described in more detail throughout the *Methods* section. Key insights include that a UAV that can survey at a GSD of 0.8 cm px^-1^ whilst at ≥ 40 m in altitude (ensuring no disturbance to animals; Pomeroy et al., 2015; Larsen & Johnston, 2023) is optimal for resolving the trade-off between data quality and the cost of data collection for species the size of UK grey seals. In addition, using a transect survey pattern instead of a radial pattern obtains volume estimates on a much wider-scale (*e.g.*, population vs small group). This new pattern allows an area of nearly 50,000 m^2^ to be surveyed in just 50 minutes, as shown in the first stage of Fig. B2.

Based on our findings, we propose our pipeline as an easily adaptable method for estimating the volume of a range of plant and animal taxa exhibiting traits similar to pinnipeds, our current study species (Table 2). For terrestrial animals, such traits include a low-to-the-ground profile, volume > 80 L, high population density, predictable geographic distribution, periods of limited mobility and inhabiting areas UAVs can fly over. Examples of animal taxa that fit these traits include larger reptiles such as crocodiles and mammals such as the hippopotamus, in addition to a range of ungulate species when resting. Many plant taxa possess all the traits that make a species well-suited to our pipeline, particularly bushes or shrubs in systems without much elevational structure such as deserts or Mediterranean habitats (Mao et al., 2021). UAVs have already been used to estimate the volume of plants such as desert shrubs (Mao et al., 2021), estimate the body length of Nile crocodiles (*Crocodylus niloticus*; Ezat et al., 2018) and assess the population sizes of ungulates (Hu et al., 2020), demonstrating these taxa are likely to be suitable for estimating volume following our pipeline. Given the widespread lack of high-quality demographic data for many of these species, such as many large reptile species (Briggs-Gonzalez et al., 2017), our pipeline presents a promising approach to collect high quality data to fill these demographic data gaps.

**Table 2.**
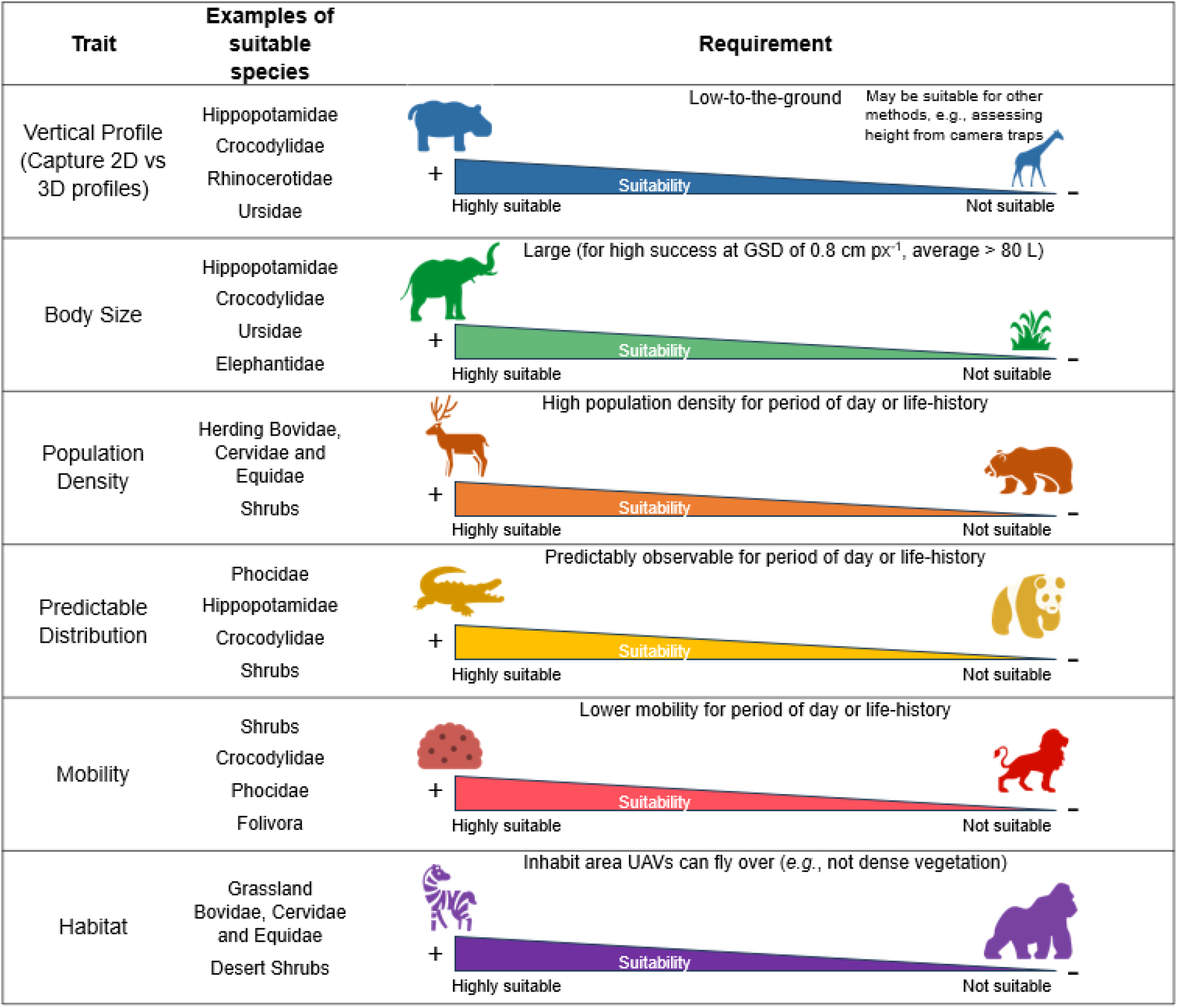
Species more suitable for volume estimates using UAVs possess several of the following six key characteristics: (1) low-to-the-ground profile, (2) volume > 100 L, (3) high population density, (4) predictable geographic distribution, (5) periods of limited mobility and (6) inhabiting areas UAVs can fly over. Here, we illustrate the adaptability of our pipeline to other settings with examples of species along the spectrum for each of these six characteristics, from more to less suitable. For example, a species must have a period during which it has low mobility so that its individuals remain stationary for the duration of a UAV survey. If an individual moves between images in a survey, the photogrammetric software will not be able to reconstruct that individual. Therefore, species such as bushes or shrubs in deserts or Mediterranean habitats would be highly suitable for estimating volume using UAVs due to their lack of movement, whereas highly mobile species such as big cats (*e.g.*, genus *Panthera*) would not.

| Trait | Examples of suitable species | Requirement |  |
| --- | --- | --- | --- |
| Vertical Profile<br>(Capture 2D vs 3D profiles) | Hippopotamidae<br>Crocodylidae<br>Rhinocerotidae<br>Ursidae | +<br>Highly suitable | Low-to-the-ground<br>May be suitable for other methods, e.g., assessing height from camera traps<br>-<br>Not suitable |
| Body Size | Hippopotamidae<br>Crocodylidae<br>Ursidae<br>Elephantidae | +<br>Highly suitable | Large (for high success at GSD of 0.8 cm px<sup>-1</sup>, average > 80 L)<br>-<br>Not suitable |
| Population Density | Herding Bovidae,<br>Cervidae and<br>Equidae<br>Shrubs | +<br>Highly suitable | High population density for period of day or life-history<br>-<br>Not suitable |
| Predictable Distribution | Phocidae<br>Hippopotamidae<br>Crocodylidae<br>Shrubs | +<br>Highly suitable | Predictably observable for period of day or life-history<br>-<br>Not suitable |
| Mobility | Shrubs<br>Crocodylidae<br>Phocidae<br>Folivora | +<br>Highly suitable | Lower mobility for period of day or life-history<br>-<br>Not suitable |
| Habitat | Grassland<br>Bovidae, Cervidae<br>and Equidae<br>Desert Shrubs | +<br>Highly suitable | Inhabit area UAVs can fly over (e.g., not dense vegetation)<br>-<br>Not suitable |

Using UAVs to estimate body size can offer advantages over traditional methods of measuring animal body size (Allan et al., 2019). Traditional methods for measuring animal body size are often time-consuming and invasive. Typically, these methods involve capturing a single individual at a time and measuring them by hand, often requiring sedatives (Manning & Goldberg, 2010; Hodgson et al., 2020; Williams et al., 2021). Using UAVs to estimate body size eliminates the need to catch or sedate animals, thus avoiding risks for the target species and the researcher (Krause et. al., 2017), and enables multiple individuals to be measured simultaneously (Shero et al., 2021). Therefore, using UAVs is faster, safer for both researchers and animals, and less invasive compared to traditional methods that use sedatives to measure body size.

Studies have estimated animal 2D body size at scale using UAVs (Gray et al., 2019; de Kock et al., 2021; Infantes et al., 2022), but recent research has shown that 3D body volume estimates are more accurate than 2D estimates (Hodgson et al., 2020; Shero et al., 2021). However, previous research only estimated the body volume of a single individual or small group of individuals at a time. Our research builds on this previous work by adopting a transect flight pattern for UAV surveys. This transect flight pattern enables the body volumes of individuals in an entire colony of taxa with similar traits to grey seals to be estimated in a single field survey. Our pipeline therefore offers a faster method to estimate body volume at the population scale. The overall mean adult body volume (133 L) we obtained is comparable to the mean body volumes estimated by UAV by Shero et al. (2021) for grey seals at Sable Island, Canada, when adjusted for the known difference in size between the Eastern and Western Atlantic grey seal populations. In addition, our recommended GSD of 0.8 cm px^-1^ is achievable with a variety of affordable, off-the-shelf UAVs at altitudes that do not disturb seals (≥ 40 m altitude; Pomeroy et al., 2015; Larsen & Johnston, 2023). As the UAVs required are off-the-shelf, relatively little training is required to operate them compared to the training required for traditional methods to estimate pinniped size involving sedative usage (*e.g.*, Hodgson et al., 2020). These practical improvements mean our pipeline is highly applicable for those responsible for managing or researching pinniped populations.

One limitation of our pipeline when surveying animal populations is that successfully estimating the volume of 100% of individuals captured in a survey is highly unlikely. Since some individuals will move between images, which prevents them from being reconstructed successfully in photogrammetric software, it is unlikely that the success rate of estimating volumes can reach 100%. For example, at our highest resolution GSD (0.8cm px^-1^), the volume of 68% of adult seals could be estimated. In these cases, the less accurate methods based on 2D photogrammetry from UAV imagery may be more appropriate, as they only require a single photo and so do not rely on the subject remaining stationary (Krause et al, 2017; de Kock et al., 2021).

A further limitation of our pipeline is that it would need to be adapted for use with aquatic species. As visibility in water is limited, the study species would have to be very close to the surface of the water. The variable visibility and movement of the water would impair the collection of different images of the same objects required for 3D photogrammetry, although novel techniques such as fluid lensing and Multispectral Imaging, Detection, and Active Reflectance (MiDAR) show this problem can be overcome in shallow waters (Chirayath & Earle, 2016; Chirayath & Li, 2019). In addition, the study species would have to have to remain stationary during surveys, in high densities, and in a predictable location for some period of time, which is unlikely in dynamic aquatic environments. However, it is possible to use UAVs to estimate the 2D body size of fully-aquatic species such as cetaceans (Gray et al., 2019), whale sharks (Whitehead et al., 2022) and corals (Levy et al., 2018). It is also possible to estimate body volume in cetaceans by creating a 3D model of the study species and scaling the model to different individuals using 2D body size estimates (Hirtle et al., 2022). Such methods present alternatives to our pipeline for some fully-aquatic species. In addition, methods exist for estimating the 3D surface area or volume of stationary aquatic species. For example, corals have been measured using handheld camera photogrammetry (Lange & Perry, 2020), satellite-based methods (Collin et al., 2021), optical coherence tomography (Jaffe et al, 2022), and underwater remotely operated vehicles (ROVs; Price et al., 2019). However, these methods are not easily adaptable to mobile species and so the challenge of developing a widely-applicable method for estimating the body volume of mobile species underwater remains unsolved.

Finding the optimal GSD at which to carry out surveys is important to resolve the trade-off between data quality and costs (Tziavou et al., 2018). Despite its importance, relatively little research has focussed on determining optimal GSDs for estimating body size, although studies have investigated the optimal GSD for wildlife detection (Frąckowiak & Goraj, 2023; Jones et al., 2023). When applying our pipeline to species of similar size to grey seals (mean volume 133 L, standard deviation 47.5 L), we recommend a maximum planned GSD of 0.8 cm px^-1^. At this GSD , the volume of 68% of adult seals (*n* = 154) could be correctly estimated, decreasing to 27% (n = 178) at 1.0 cm px^-1^. This 25% decrease in GSD therefore led to a 41% decrease in successful volume estimates-a greater decrease than we expected. The percentage of successful volume estimates was similar at 1.0 cm px^-1^ and 1.2 cm px^-1^ (28%, *n* = 87), indicating a plateau in feasibility above 1.0 cm px^-1^. It is possible the larger-than-expected decrease in success rate and subsequent plateau is due to the existence of a threshold value between GSDs of 0.8 cm px^-1^ and 1.0 cm px^-1^ at which images are no longer high enough resolution for Pix4D to accurately reconstruct seal-sized objects. For animals larger than grey seals, surveying at a GSD of 0.8 cm px^-1^ will give high enough quality data to obtain accurate volume estimates at scale, but further research may determine that surveying at a higher GSD is also possible. We recommend further research on the optimal GSD for species that are smaller in size than grey seals as lower GSDs are likely to be required for smaller species.

Our optimal GSD of 0.8 cm px^-1^ is higher than the average GSD used in previous work estimating body volume of pinnipeds using UAVs (Hodgson et al. 2020; Shero et al., 2021). Therefore, we demonstrate that a wider range of UAVs or altitudes can be used for this research than have been used previously. We expect GSDs less than 0.8cm px^-1^ would further improve the percentage success rate of reconstructing seals so their volumes can be estimated and the accuracy of these estimates. However, we were limited to surveying at an altitude ≥ 40 m to ensure seals were not disturbed (Pomeroy et al., 2015; Larsen & Johnston, 2023) and by the resolution of off-the-shelf, affordable UAVs.

The population-scale, continuous body size data collected following the pipeline presented here have a wide range of uses. One key application is using a time series of such data to parameterise demographic models such as integral projection models (IPMs; Easterling et al., 2000; Merow et al., 2014). Such demographic models can be used to better understand how population size and structure is changing through time, and may change into the future and in response to key threats like climate change or habitat degradation (Gerber & Heppell, 2004). These data can be integrated into an IPM thanks to inverse-modelling techniques (González et al., 2016), even for species with high mobility like pinnipeds (Wielgus et al., 2008), and thus where no longer observing an individual in the next census may not imply mortality. To the best of our knowledge, though, no IPMs have been constructed for pinniped species using body size data. Using body size data, body condition indices can be developed (Krause et al., 2017). Such indices could take into account other data that can be obtained from UAV imagery. For example, thermal imagery could be used to give a score of animal health or reproductive ability (Jeelani & Jeelani, 2019), given the importance of metabolic rate to key processes such as survival and reproduction (Brown et al., 2004; Savage et al., 2004). A continuous body condition index could also be used to parameterise demographic models like IPMs. In addition to parameterising demographic models, data obtained via our pipeline can facilitate research into tracking energy-flow dynamics (Shero et al., 2021), assessing individual health (Hodgson et al., 2020) and fitness (Stevenson & Woods, 2006) across populations, investigating variation in metabolic activity (Sparling et al., 2006), comparing health of different populations (Sweeney et al., 2014), and detecting how environmental conditions affect body size (Berger, 2012).

Currently, bottlenecks exist in our pipeline at the seal identification and volume estimation stages of the pipeline, which are performed manually. To date, machine learning has been used to successfully automate or semi-automate the process of estimating 2D body size or identifying animals from images, for example, in cetaceans (Gray et al., 2019) and pinnipeds (Infantes et al., 2022). Therefore, we expect future work taking advantage of machine learning technology will be able to overcome these bottlenecks in our pipeline to greatly decrease the time taken to identify and estimate the volume of individual seals.

### Conclusions

Our pipeline enables accurate body volume to be estimated in a time-and cost-effective manner using off-the-shelf, multirotor UAVs whilst minimising the disturbance caused to the study species. The ability to obtain accurate body volume estimates of taxa that are otherwise challenging to study at scale enables the development of continuous, size-or body condition-structured demographic models. This work therefore opens up gaps in the Tree of Life (Lebreton et al., 2012; Salguero-Gómez et al., 2015; Salguero-Gómez et al., 2016) to demographic studies that will improve our ability to protect and conserve species into the future.

## Acknowledgments

We acknowledge support from 10% for the ocean, the Eurofins Foundation Fund and Brasenose College, University of Oxford. We thank the Isles of Scilly Wildlife Trust and National Trust for their assistance with data collection. We thank R. Salguero-Gómez and M. Burns Zaragoza for their invaluable comments to improve this manuscript, M. Chopra, L. Beaupere and R. Salguero-Gómez for help with fieldwork, and M. Stone for his assistance with validation of the accuracy of volume estimates.

## Footnotes

1 Undertaking censuses of any wild animal population with UAVs requires trained operators as well as obtaining permissions and carrying-out pre-site assessments (Barnas et al., 2020). In the UK, UAV operators require a Civil Aviation Authority A2CofC Certification, which we obtained prior to our flight missions. We obtained permissions from local land managers for the Isles of Scilly and Farne Islands, and further permissions from local airspace controllers given the proximity of an airport to the Isles of Scilly field sites.

## Boxes, Tables and Figures

### Box 1

Ground sampling distance for estimating volume

Ground sampling distance (GSD) is a measure of the resolution of an image, representing the true distance between the centre of two adjacent pixels. A higher GSD, therefore, results in a lower resolution image. For example, if a camera’s minimum GSD at a certain altitude is 0.8 cm pixel^-1^, the smallest details that could be obtained from an image taken at said altitude would be 0.8 cm measured linearly (Fig. B1). When planning surveys, GSD is dependent on flight altitude and three camera parameters: sensor width, focal length, and image width. Because the GSD can be calculated for any UAV camera at a given altitude, once the maximum GSD that enables many accurate body volume estimates of an animal species to be obtained, one can determine whether any UAV can estimate the body volume of animals of that size at the altitude required to avoid disturbance. GSD is calculated as:

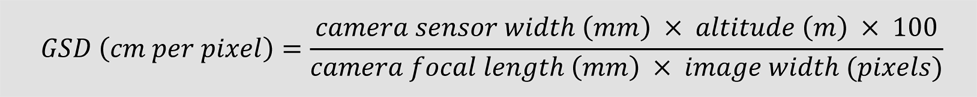

For example, the DJI Mavic 2 Enterprise Advanced has a visual-range camera with a 1/2” CMOS sensor, which has a sensor width of 6.4 mm, a focal length of 4mm, and an image width of 8,000 pixels. Therefore, taking images at 40 m altitude gives a GSD of 0.8 cm px^-^

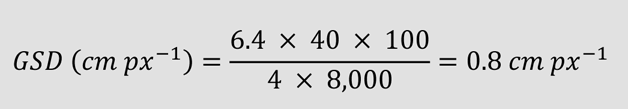

**Figure B1.**
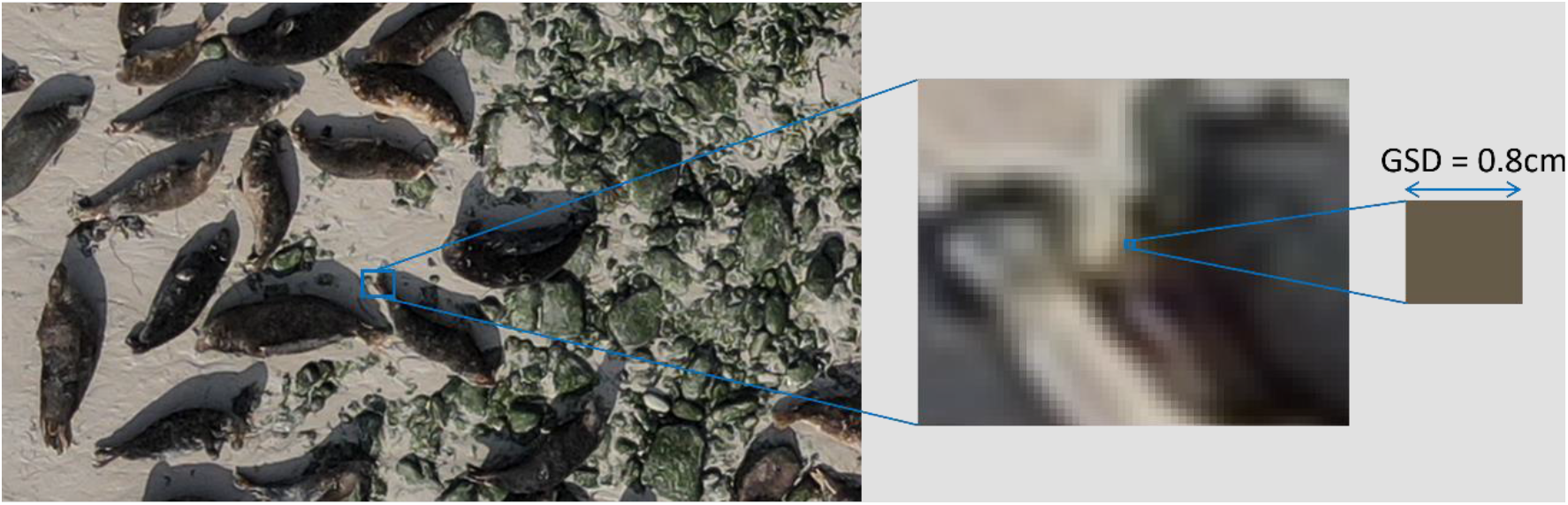
Ground sampling distance (GSD) is a measure of the resolution of an image, representing the true distance between the centre of two adjacent pixels. Therefore, if each pixel represents 0.8 cm in real life, the GSD of the image is 0.8 cm px-1. A GSD of 0.8 cm px-1 is illustrated here, width the width of each pixel representing 0.8 cm in real life.

### Box 2

Pipeline and best practices

Our pipeline has five key stages (Fig. B2), starting with the pre-fieldwork stage to ensure the appropriate training and planning have been carried out. The second stage describes best practises in the field. When in the field, UAV batteries are a key limiting factor due to cost and transportability issues (*e.g.*, one DJI Mavic 2 Enterprise Advanced battery is £160), as sufficient batteries must be taken into the field for the day’s activities. Therefore, optimising the number of UAV batteries required in the field is important. In the field, keeping a flight log detailing survey flight times, locations, and SD card used makes post-field processing easier. The third stage is the pre-processing stage, which helps to make the subsequent processing stage easier. When undergoing data pre-processing, removing blurry images or images where a large number of seals were moving can allow 3D models to be reconstructed where reconstruction would otherwise be impossible. In stage four, the processing stage, images are stitched together to form a geometrically accurate 3D model, seals are identified and counted, and individual volumes are estimated. Exporting the orthomosaic into manual counting software such as ‘DotDotGoose’ (v1.5.3; Ersts, 2022) helps to accelerate the seal identification and counting step. Finally, in stage five, the resulting body volume estimates can be used for their intended purpose.

**Figure B2.**
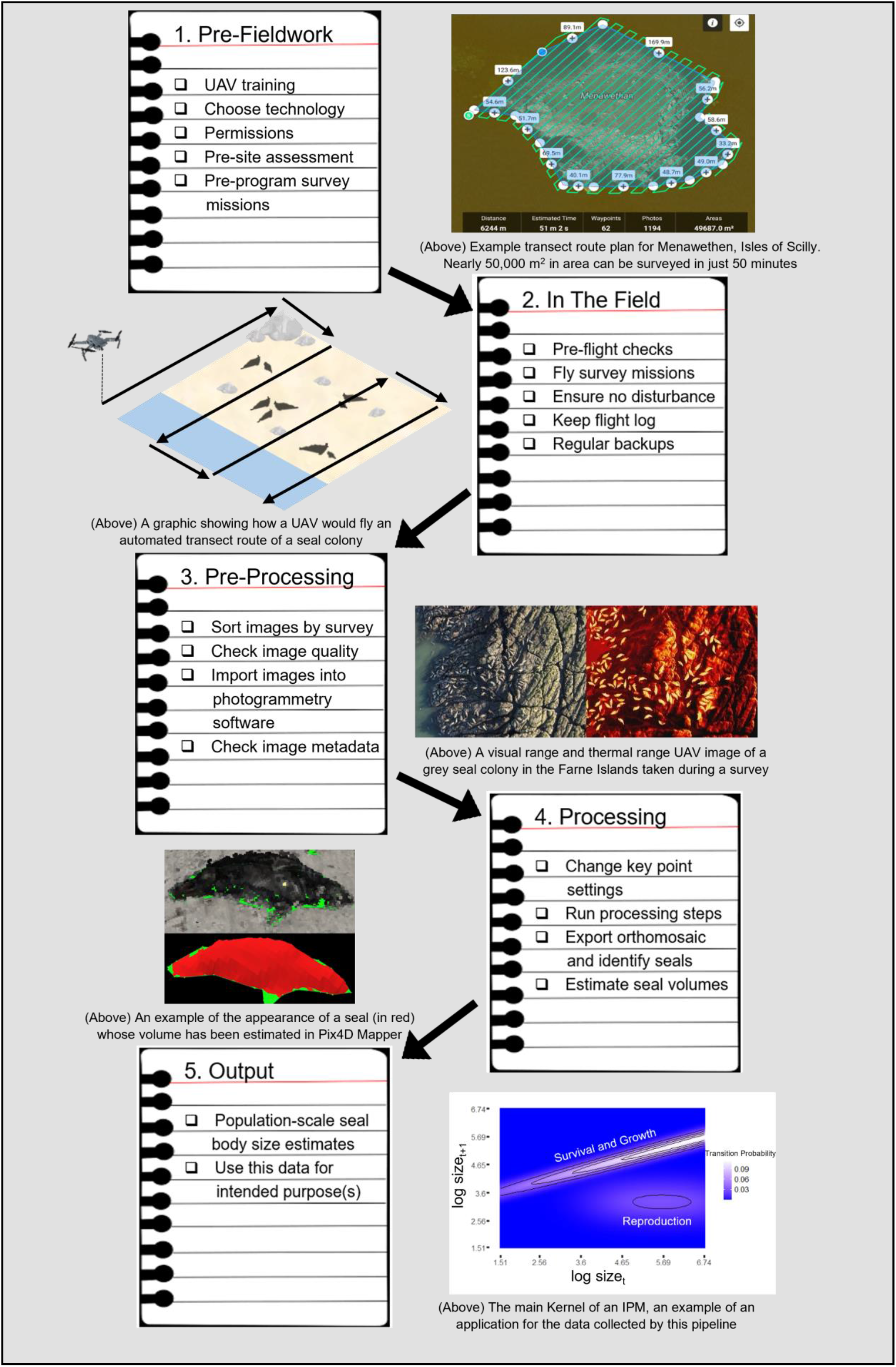
Overview of the UAV pipeline we developed to estimate the body volume of pinnipeds at scale, from pre-fieldwork to output stages.

## Appendix A

Table A1 gives a breakdown for how the values for total time required were calculated. Depending on conditions, a drone battery change is required for approximately every 22 minutes of survey time when using the DJI Mavic 2 Pro or DJI Mavic 2 Enterprise Advanced UAVs. This is a conservative estimate to ensure plenty of battery (return at 30 - 40%) to return to and land on the boat, and adds an additional constraint as the number of batteries you can take into the field is limited by their cost and space for equipment. Each battery change takes approximately 5 - 10 minutes of flying back, landing, changing the battery, taking off and returning to the point the survey was paused. For the Isles of Scilly, carrying out the census at 40 m therefore adds an additional 490 minutes and 16 battery changes compared to censusing at 60 m, the equivalent of approximately two days in the field.

**Table A1.** The time for surveys increases with decreasing ground sampling distance (GSD; Box 1). A breakdown of the time it would take to census the Isles of Scilly grey seal population at different GSDs using the DJI Mavic 2 Enterprise Advanced UAV.

| **GSD** | **0.8cm px<sup>-1</sup>** | **1.0cm px<sup>-1</sup>** | **1.2cm px<sup>-1</sup>** |
| --- | --- | --- | --- |
| Flight Time<br>(Minutes) | 1263 | 1056 | 934 |
| Battery Changes<br>Required | 58 | 48 | 43 |
| Battery Change<br>Time (Minutes) | 580 | 480 | 430 |
| Total Time Required<br>(Minutes) | 1843 | 1536 | 1364 |

## Notes

### Competing Interest Statement

The authors have declared no competing interest.

